# Tree diversity intensifies soil microorganism-tree interactions

**DOI:** 10.64898/2026.05.05.722867

**Authors:** Haikuo Zhang, Naili Zhang, Helge Bruelheide, Xiaojuan Liu, Shan Li, Zhuo Yang, Yanjiang Cai, Alexandra M. Klein, Steffen Seitz, Thomas Scholten, Yvonne Oelmann

**Affiliations:** Geoecology, Department of Geosciences, University of Tübingen, 72070 Tübingen, Germany; Soil Science & Geomorphology, Department of Geosciences, University of Tübingen, 72070 Tübingen, Germany; State Key Laboratory of Efficient Production of Forest Resources, Key Laboratory for Silviculture and Conservation of the Ministry of Education, College of Forestry, Beijing Forestry University, 100083 Beijing, China; Institute of Biology/Geobotany and Botanical Garden, Martin Luther University Halle-Wittenberg, 06108 Halle, Germany; German Centre for Integrative Biodiversity Research (iDiv) Halle-Jena-Leipzig, 04103 Leipzig, Germany; State Key Laboratory of Forage Breeding-by-Design and Utilization, Institute of Botany, Key Laboratory of Vegetation and Environmental Change, Institute of Botany, Chinese Academy of Sciences, 100093 Beijing, China; University of Chinese Academy of Sciences, 100049 Beijing, China; National Key Laboratory for Development and Utilization of Forest Food Resources, College of Environment and Resource Sciences, College of Carbon Neutrality, Zhejiang A&F University, 311300 Hangzhou, China; Nature Conservation and Landscape Ecology, University of Freiburg, 79106 Freiburg, Germany; Physical Geography, Institute of Geography, Osnabrück University, 49074 Osnabrück, Germany

## Abstract

- A productivity-driven higher nutrient demand of trees in diverse mixtures is frequently reported. Yet, it remains unclear how tree diversity influences microorganisms-plants interactions, in which microbes facilitate tree nutrient acquisition in exchange for carbon (C) to meet the resource demand of both.
- Using a long-term tree diversity experiment in the subtropics, we assessed microbial investment in C-, nitrogen (N)-, and phosphorus (P)-acquiring enzymes in litter and mineral soil, testing the effects of tree species richness and mycorrhizal type (arbuscular (AM)- vs. ectomycorrhizal (EcM)-associated tree species).
- With increasing tree species richness, microbial investment in C acquisition decreased, while investment in N and/or P acquisition increased in litter and in mineral soil. In mineral soil of AM-associated tree mixtures, ecoenzymatic stoichiometry revealed a shift from microbial investment in C toward P acquisition as tree species richness increased.
- Our findings suggest that tree diversity strengthens microbe-tree interactions in terms of C-for-nutrient exchange. This highlights the key role of soil microorganisms, particularly in AM symbiosis, shaping tree diversity-biogeochemical feedbacks.

## Introduction

Biodiversity enhances ecosystem’s primary productivity (Scherber *et al*., 2010; Isbell *et al*., 2017; Huang *et al*., 2018; Wagg *et al*., 2022). Increased biomass production in diverse plant communities is accompanied by higher demand for nutrients (Cardinale *et al*., 2007), e.g., nitrogen (N) and phosphorus (P). Microorganisms in litter and mineral soil play a crucial role in plant productivity (Griffiths *et al*., 2021) by regulating nutrient supply (Van Der Heijden *et al*., 2008; Čapek *et al*., 2018). They mineralize organic matter (OM) into plant-available inorganic nutrients (Van Der Heijden *et al*., 2008), facilitated by microbially secreted extracellular enzymes (e.g., proteases and chitinases that degrade N-containing complex proteins and chitin into bioavailable amino acids) (Das & Varma, 2011; Nannipieri *et al*., 2012; Burns *et al*., 2013; Nannipieri *et al*., 2018). In turn, plants fuel microbial activities through the transfer of assimilated C, creating a feedback loop where microbes may reduce the production of energy-intensive C-acquiring enzymes (i.e., extracellular enzymes such as cellulases that release assimilable C from complex OM) to follow the principle of maximizing energy gains (Schnitzer *et al*., 2011; Jacoby *et al*., 2017). This mutualistic interaction between microorganisms and plants may be enhanced in a diverse plant community, where photosynthates, varied plant root exudates, and litter provide abundant C inputs (Steinauer *et al*., 2016; Eisenhauer *et al*., 2017; Huang *et al*., 2018). Such inputs may allow microbes to downscale costly C-acquisition and redirect resources toward enzymes mobilizing limiting nutrients like N and P, thereby enhancing plant growth and nutrient cycling in more diverse mixtures (Schnitzer *et al*., 2011). Evidence from a long-term plant diversity experiment in a grassland ecosystem and a recent global meta-analysis confirms that plant diversity promotes P acquisition by stimulating microbial enzyme activities (Oelmann *et al*., 2011; Hacker *et al*., 2015; Chen *et al*., 2022). However, whether the increased microbial investment in nutrient acquisition is linked to a decrease in the activity of C-acquiring enzymes remains unresolved. Elucidating this potential trade-off between the microbial investment in nutrient- and C-acquisition is essential for fully understanding the biodiversity-ecosystem functioning relationships.

Mycorrhizal fungi can directly supply essential nutrients to host plants (Martin *et al*., 2008; Van Der Heijden *et al*., 2008; Pritsch & Garbaye, 2011; Lambers *et al*., 2018). By forming symbiotic associations with plant roots, these fungi provide plants with a range of limiting nutrients via their extensive hyphal networks in exchange for plant-derived photosynthates (e.g., carbohydrates and lipids) (Smith & Read, 2010; Smith & Smith, 2011). In forests, the most prominent and frequently studied groups of mycorrhizal fungi are arbuscular mycorrhizal (AM) and ectomycorrhizal (EcM) fungi (Smith & Read, 2010; Tedersoo & Bahram, 2019). AM and EcM fungi form distinct symbiotic structures and colonize tree roots spanning from the litter layer to mineral soil (Martin *et al*., 2008; Smith & Read, 2010; Tedersoo *et al*., 2020; Freschet *et al*., 2021). In AM symbioses, fungi form intimate intracellular associations with root cells via arbuscules, facilitating efficient C-nutrient exchange (Smith & Read, 2010). The AM fungi generally have a limited repertoire of extracellular enzymes targeting simple organic compounds. For example, they contribute little directly to total soil phosphatase activity but often enhance phosphatases derived from plant roots and free-living microbes, thereby promoting the acquisition of organic P (Rosling *et al*., 2016; Jiang *et al*., 2021; Wang *et al*., 2023). In contrast, EcM fungi develop a mantle surrounding the root tip, a structure that supports more autonomous extracellular enzyme secretion and nutrient acquisition (Martin *et al*., 2008). Furthermore, EcM fungi are capable of producing a broader spectrum of extracellular enzymes, including peroxidases, laccases, proteases, and phosphatases (Talbot *et al*., 2013), enabling them to degrade complex soil organic matter (SOM) such as lignin, cellulose, and proteins (Lindahl & Tunlid, 2015). A large-scale tree diversity experiment in the subtropics demonstrated that AM and EcM host trees differentially modulated the biodiversity–productivity relationship largely through their contrasting nutrient acquisition strategies (Deng *et al*., 2023). Specifically, compared with EcM-associated trees, AM-associated trees exhibited higher-quality, more decomposable litter, higher leaf nutrient resorption efficiency, and faster litter decomposition rates in more diverse mixtures, thereby maintaining a productivity advantage (Steidinger *et al*., 2019; Deng *et al*., 2023). Such conditions imply that increased tree diversity may enhance interactions between AM fungi and free-living microbes to mobilize inorganic nutrients, such as P, from high-quality OM in a way that exceeds the autonomous EcM enzyme secretion strategy (Rosling *et al*., 2016; Jiang *et al*., 2021). Simultaneously, the AM fungi/free-living microbe interaction might lessen the need to invest in C-acquiring enzymes because of the combination of plant-derived C transfer by photosynthates (AM fungi) and roots exudates (free-living microbes). In line, functional predictions of soil microbial communities in the same tree diversity experiment further indicated that microbial subcommunities in AM-associated tree mixtures are more strongly linked to P-cycling enzyme than those in EcM-associated tree mixtures, and also more strongly than to C-cycling enzyme (Singavarapu *et al*., 2023). However, whether these mycorrhiza-specific differences in the genomic potential of microbial communities translate into real functioning remains unclear, particularly given that the differences disappeared at higher tree diversity (Singavarapu *et al*., 2023), which might not apply to enzyme synthesis and consequently enzyme activities.

To better understand the roles of soil microorganisms in providing and receiving responses to tree diversity and the mycorrhizal types of host plants, we employed an ecoenzymatic stoichiometry approach. To this end, we assessed the microbial investment in N, P, and C acquisition based on the N-acquiring (β-1,4-N-acetylglucosaminidase, NAG; leucine aminopeptidase, LAP), P-acquiring (acid phosphatase, ACP), and C-acquiring (β-1,4-glucosidase, BG) enzyme activities in litter and mineral soil. The individual activities were then stoichiometrically combined to contrast the microbial investment into nutrient versus C acquisition. We relied on a large forest tree diversity experiment in subtropical China (BEF-China platform), ranging from monocultures to mixtures with 16 tree species.

We hypothesize that:

□ Soil and litter microorganisms shift their enzymatic investment towards N and P acquisition at the expense of C acquisition under high tree species richness.
□ This shift in microbial nutrient investment varies with the mycorrhizal type of the host trees, with a more pronounced shift toward P acquisition in AM-associated tree mixtures.

## Materials and Methods

### Study area and experimental design

The study was conducted at two main experimental sites (i.e., Site A and Site B) of the Biodiversity–Ecosystem Functioning Experiment China (BEF-China, https://bef-china.com) platform, located in Xingangshan, Dexing, Jiangxi Province, China (29.08–29.11° N, 117.90–117.93 ° E). This region exhibits a subtropical monsoon climate with 16.7 °C mean annual temperature and 1821 mm mean annual precipitation (Yang *et al*., 2013). The soil types are mainly classified as Cambisols with Anthrosols in downslope positions and Gleysols in valleys (Seitz *et al*., 2016; Scholten *et al*., 2017). Soils of both experimental sites are loamy with silt contents between 44% and 50%, a mean bulk soil density of 1.3 g cm^-3^, and acidic with values (pH_KCl_) from 3.2 to 4.7. Soil organic C varies between 4.9%–2.7% and total N between 0.5%–0.2% in the A horizon. The CEC is similar at both experimental sites. In total, 566 plots (with a size of 25.8 m × 25.8 m) were planted in site A (18.4 ha, 271 plots) in 2009 and site B (20 ha, 295 plots) in 2010 (Bruelheide *et al*., 2014). Each plot has 400 tree individuals at equal distances of 1.29 m. Tree species richness ranges from monocultures to mixtures of 2, 4, 8, 16, and 24 species. Species compositions of the different diversity levels were based on 40 local tree species, including AM/EcM host trees, and realized in random and trait-informed extinction scenarios (see tree species list in Table S9). The random extinction scenarios were constructed by a broken stick design, starting from three different but overlapping sets of 16 species per site.

### Soil and litter sampling

The soil sampling was conducted in October–December 2023 and consisted of 293 plots, containing 146 plots selected on Site A and 147 on Site B (including 1-, 2-, 4-, 8-, and 16-species mixtures; excluding shrub, conifer, free succession, and genetic plots from the original BEF-China setup). In each plot, after removing the surface organic layer, a total of nine soil cores were obtained to a depth of 5□cm and mixed to form a composite soil sample (Scholten *et al*., 2017). The soil samples were immediately transported to the laboratory, then sieved (2 mm mesh) to remove visible plant debris and soil fauna residues, and stored at -20 □ until the subsequent soil enzyme activity assay.

To provide a comparative perspective on decomposer nutrient dynamics in the litter layer, we re-analyzed previously published litter enzyme data from a subset of 105 of these plots. It is important to note that the litter samples used in this study were collected by Zhang *et al*. (2018) in October 2014. The samples were collected on 105 plots, containing 49 plots selected on Site A and 56 on Site B (including 1-, 2-, 4-, 8-, and 16-species mixtures; from the “very intensively studied plots” of the original BEF-China setup). The mixed leaf litter was collected from five randomly selected positions in each plot. More details on the litter sampling process can be found in Zhang *et al*. (2018).

### Ecoenzymatic activity assay

Both soil and litter enzyme activities were measured by the microplate fluorometric techniques (German *et al*., 2011; Zhang *et al*., 2018; Zhang *et al*., 2022). For the soil enzymes (BG, NAG, LAP, and ACP) activity assay, all soil samples were transferred from a -20 □ freezer to a 4 □ refrigerator until thawed before measuring. The 1.5 g thawed fresh soil was homogenized in 125 mL 50 mM sodium acetate buffer (pH = 5.0) in a magnetic blender for 1 min. The 200 µL soil slurry was dispensed into the 96-well microplate, and each sample included three columns with 8 replicate wells. The first column was a 200 µL soil slurry + 50 µL sodium acetate buffer (50 mM), the second column was a 200 µL soil slurry + 50 µL standard solution (10 µΜ of 4-methylumbelliferone or 7-amino-4-methylcoumarin), and the third column was a 200 µL soil slurry + 50 µL specific substrate solution (4-MUB-β-D-glucopyranoside for BG; 4-MUB-N-acetyl-β-D-glucosaminide for NAG; L-leucine-7-amido-4-methylcoumarin hydrochloride for LAP; 4-MUB phosphate for ACP). The microplate was incubated in the dark at 25 °C for 3 h. The fluorescence value was determined using a microplate reader (Biotek^®^ Synergy H1, Winooski, VT, USA) with 365 nm excitation and 450 nm emission. Ecoenzymatic activities were expressed as nmol g^-1^ dry soil h^-1^.

The litter enzymes (BG, NAG, and ACP) activity was determined by Zhang et al. immediately after sampling in 2014 (Zhang *et al*., 2018). Except for the weight of the sample, the methods of litter enzyme activity assay are the same as those for soil. More details on the experimental process can be found in Zhang *et al*. (2018).

### Ecoenzymatic stoichiometry

To explore the extent of soil and litter microbial investment into C and nutrient acquisition, the vector analysis (including vector angle and vector length) of ecoenzymatic stoichiometry was calculated using the following equations (Moorhead *et al*., 2016):

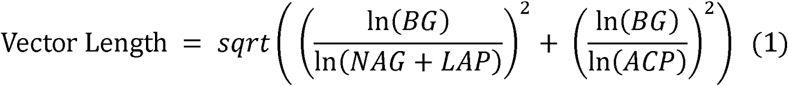

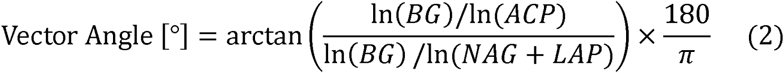

where the BG, NAG+LAP, and ACP represent the corresponding ecoenzymatic activities. The vector angle < 45° and > 45° represent the relative degree of investment into N and P acquisition of microbial communities, respectively. A vector angle of < 45° represents greater microbial investment into N than into P acquisition. Conversely, a vector angle of > 45° indicates greater microbial investment into P than into N acquisition. The vector length represents the relative degree of microbial investment into C acquisition: the longer the vector length, the greater is the investment into C relative to the nutrient acquisition.

### Statistical analyses

Linear mixed-effects models (LMEs) were used to evaluate the effect of tree species richness (1-, 2-, 4-, 8-, and 16-species mixtures), tree mycorrhizal type (AM, EcM, and AM + EcM), tree species richness & mycorrhizal type interactions, and topographic factors (altitude, aspect, and slope) on soil & litter ecoenzymatic activities, vector length, and vector angle, with the sampling site as a random factor. To test for overall differences between mycorrhizal groups (AM, EcM, and mixed AM + EcM), we additionally performed one-way ANOVA followed by Tukey’s Honestly Significant Difference (HSD) tests (n = 93, 67, and 133 for soil; n = 59, 25, and 21 for litter, respectively). This complementary analysis was designed to illustrate the main effects of mycorrhizal type independent of tree species richness, whereas the effects of tree species richness and its interaction with mycorrhizal type were examined using the LMEs described above. We also used LMEs to analyze ecoenzymatic activities, vector lengths, and vector angles in plots exclusively containing AM, EcM, or both host tree types (n = 91, 67, and 64 for soil, respectively; n = 57, 25, and 13 for litter, respectively) across a gradient of 1-, 2-, and 4-species mixtures, with the tree species richness and topographic factors as fixed factors and sampling site (site A and site B) as a random factor. The constrained tree species richness level is due to the fact that none of the more diverse 8- and 16-species mixtures exclusively contains either AM or EcM host trees. The assignment of the mycorrhizal type matches the observed microorganism community in soil. The above statistical analyses were performed using the “stats” and “lme4” packages of the R software (version 4.4.0, http://www.R-project.org).

## Results

Higher tree species richness significantly decreased C-acquiring enzyme activities (Fig. 1a), while it did not affect N- and P-acquiring enzyme activities in litter (Figs. 1, c and e). Conversely, tree species richness significantly increased the N- and P-acquiring enzyme activities (Figs. 1, d and f), but it had no effect on C-acquiring enzyme activities in mineral soil (Fig. 1b). The vector angle greater than 45° indicated more investment in P acquisition than in N acquisition in both litter and mineral soil (Figs. 2, c and d). In the litter, vector angles and tree species richness were positively related (Fig. 2c), indicating greater allocation to P acquisition with increasing tree species richness. Both for litter and soil, tree species richness significantly decreased the vector lengths, and thus, the microbial investment in C acquisition compared to N and P acquisitions (Figs. 2, a and b).

**Fig. 1.**
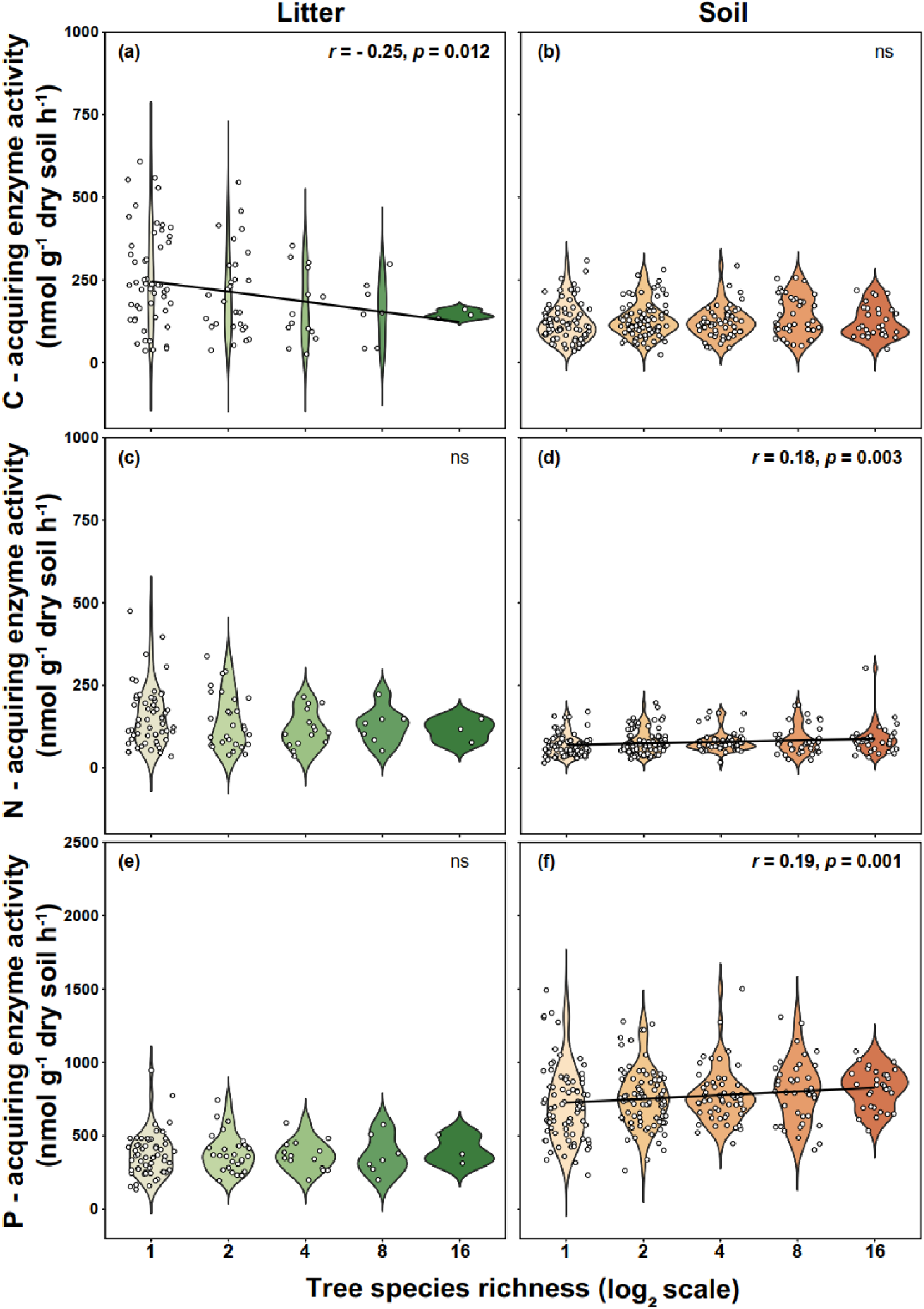
Variations in litter (a, c, and e) and soil (b, d, and f) C-, N-, and P-acquiring enzyme activities with tree species richness. Black regression lines are based on linear mixed-effects models (predicted means).

**Fig. 2.**
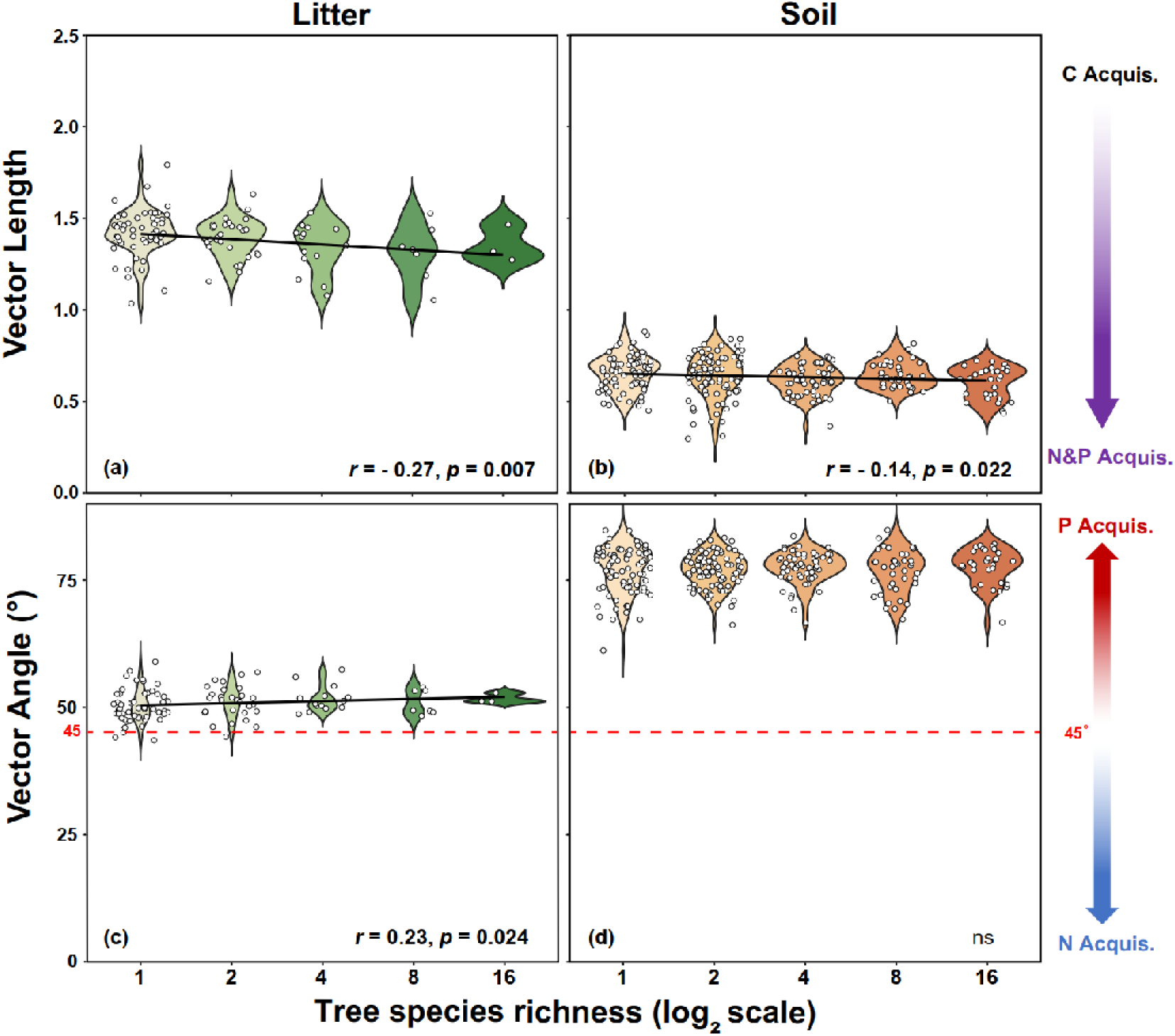
Variations in litter and soil ecoenzymatic stoichiometry with tree species richness. Subgraphs (a) to (d) show the changes in litter and soil microbial C vs. nutrients acquisition (Vector Length) and microbial N vs. P acquisition (Vector Angle) in relation to tree species richness. Black regression lines in subgraphs (a) to (c) are based on linear mixed-effects models (predicted means).

The mycorrhizal type of the host trees in mixtures, both alone and in interaction with tree species richness, significantly influenced enzyme activities and stoichiometry, but this effect was confined to the mineral soil (Figs. 3 and 4 compared to Figs. S1 and S2). If AM-associated trees were present in mixtures (AM, AM+EcM), P-acquiring enzyme activities in soil were higher than in mixtures without AM-associated trees (EcM-associated, Fig. 3c). Similarly, the vector angles in soil of the mixtures with AM-associated trees were higher, indicating more investment in P acquisition than in those with EcM-associated trees (Fig. 3e).

**Fig. 3.**
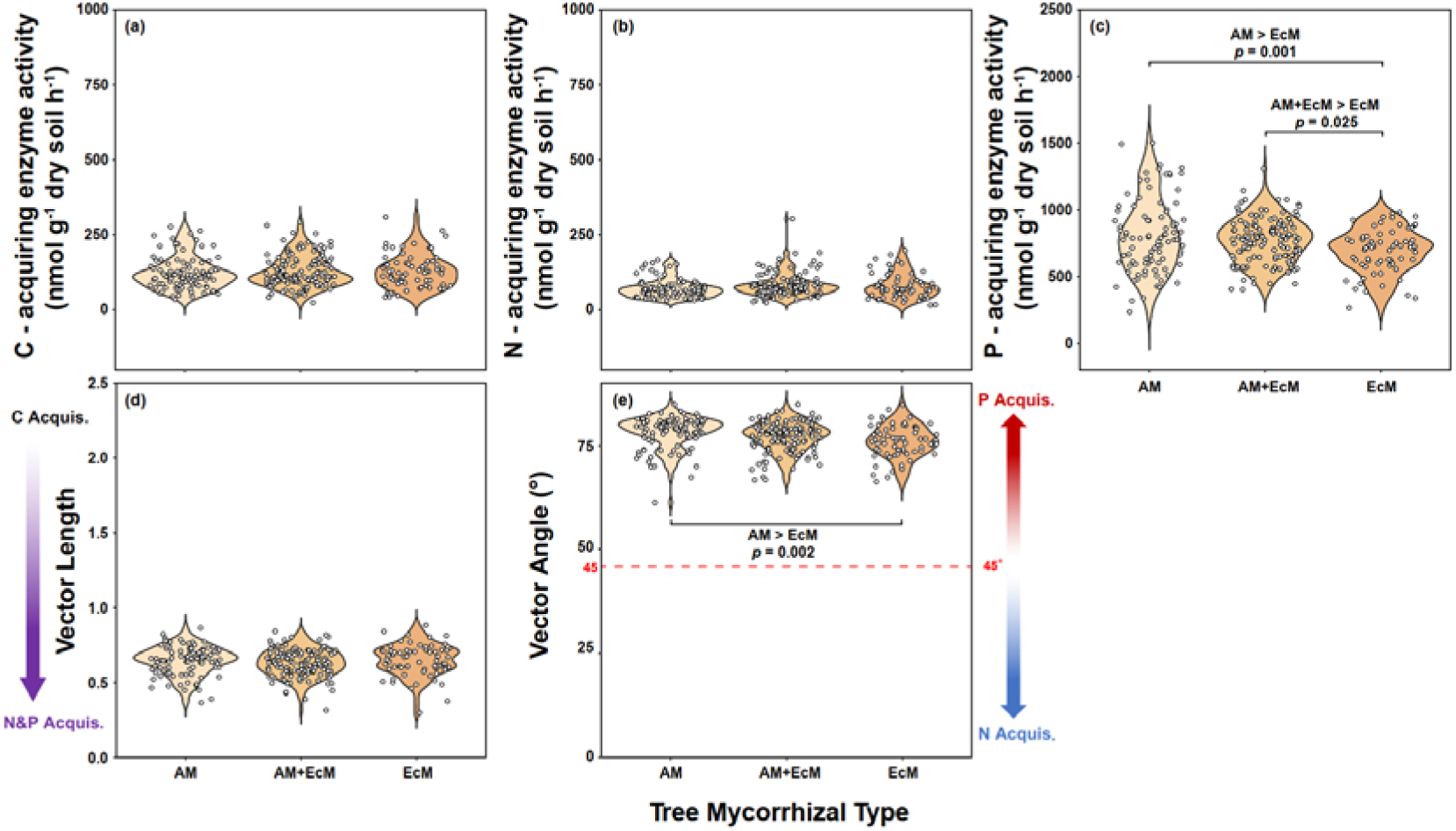
Soil enzyme activities (a-c) and microbial C vs. nutrient acquisition (d, e) in relation to tree mycorrhizal types for plots with mixtures containing only arbuscular mycorrhizal (AM) host tree species, only ectomycorrhizal (EcM) host tree species, and both mycorrhizal types (AM + EcM).

**Fig. 4.**
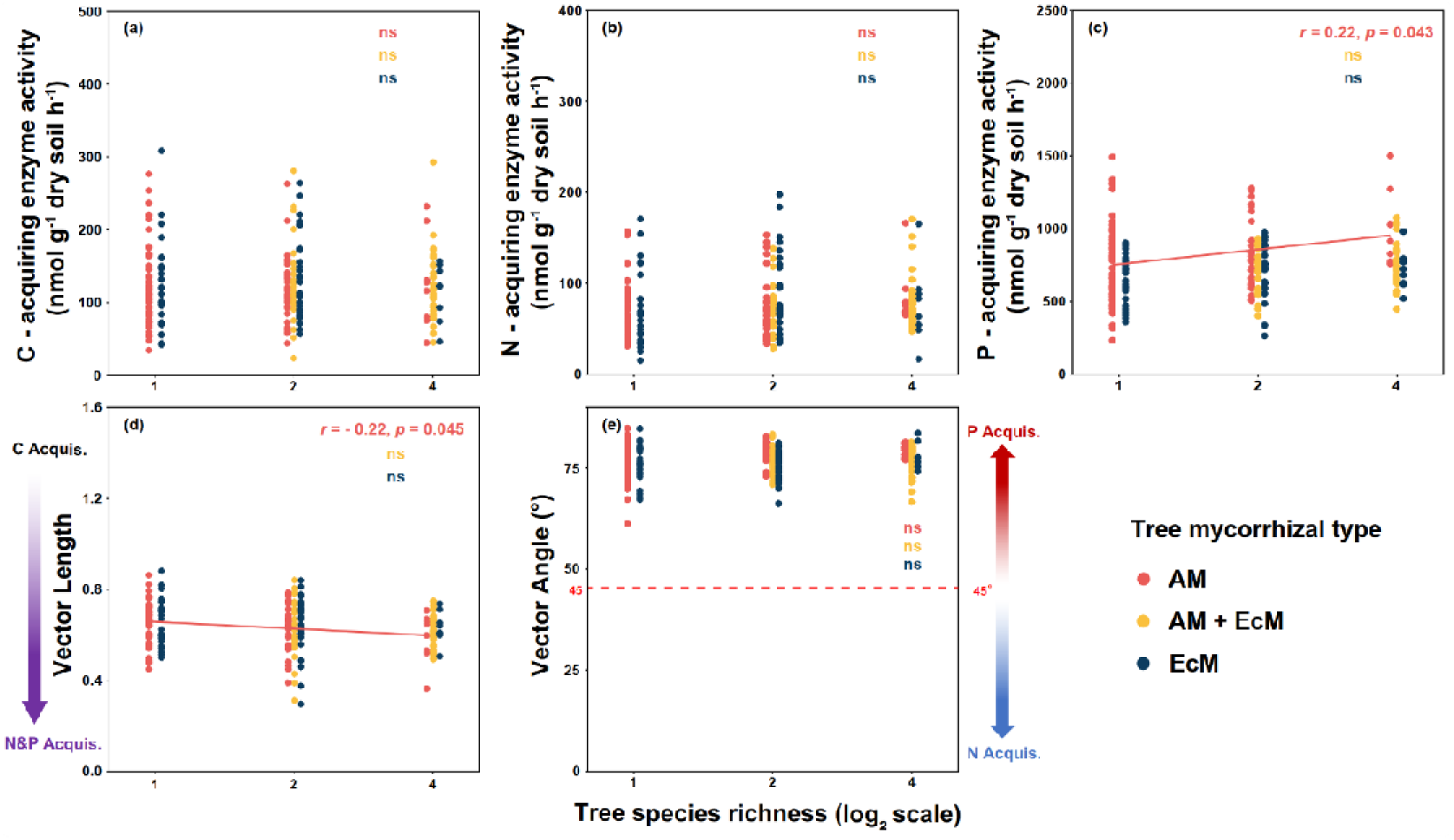
Soil enzyme activities (a-c) and microbial C vs. nutrient acquisition (d,e) in relation to tree species richness (one, two, four) for plots with mixtures containing only arbuscular mycorrhizal (AM) host tree species, only ectomycorrhizal (EcM) host tree species, and both tree species types (AM + EcM). Each dot represents a plot, and colors indicate different tree mycorrhizal types. Regression lines are based on linear mixed-effects models (predicted means). Solid lines indicate statistically significant relationships (*p* < 0.05).

With increasing tree species richness, P-acquiring enzyme activities in soil increased in mixtures with only AM-associated trees (Fig. 4c). For this evaluation, we had to exclude the 8- and 16-species mixtures, which did not contain compositions without AM-associated tree species. Similar to the overarching negative effect of tree species richness on investment into C acquisition described above, the vector lengths of soil in mixtures with AM-associated trees decreased with increasing tree species richness (Fig. 4d).

## Discussion

### Tree species richness enhances microbial nutrient acquisition

We found that higher tree species richness promoted the investment of soil microorganisms into N and/or P acquisition in litter and mineral soil (Figs. 1, d and f; 2, c). It has been demonstrated that higher tree species richness is associated with higher biomass production, which results in more nutrient demand for tree growth (Deng *et al*., 2023). This productivity-driven nutrient demand is not only met by nutrient resorption by trees before leaf shedding, but also requires a large amount of nutrients to be taken up from soil (Deng *et al*., 2023). In the latter case, the trees benefit from microbially derived nutrients (Jacoby *et al*., 2017; Semchenko *et al*., 2022). Specifically, the main explanation for increased microbial investment in N and P acquisition by tree species richness could be the increased availability of substrates for soil microorganisms: On the one hand, SOM serves as the substrate for enzymatic degradation. For our study site, an increase in SOM by tree species richness was reported and could enhance investment into nutrient acquisition by soil microorganisms (Hacker *et al*., 2015; Li *et al*., 2019). On the other hand, previous studies demonstrated that plant mixtures provided more diverse litter (De Groote *et al*., 2018; Hoeber *et al*., 2020), which increased N and P sources for soil microorganisms. The inference was also confirmed by a recent global synthesis that soil net N mineralization rates are promoted in the species-rich plant mixtures (Chen *et al*., 2021), a pattern attributed to SOM accumulation and enhanced microbial activity driven by higher-diverse litter inputs from increased plant diversity. Similarly, another recent global comprehensive study showed that highly diverse plant species mixtures supplemented soil bioavailable P by promoting soil P-acquiring enzyme activity (Chen *et al*., 2022).

The microbial investment into P acquisition in litter and soil played a more important role than that related to C and N acquisition (Figs. 1 and 2). The shortage of P resources in soil and the increased nutrient demand of plants and soil microorganisms in our study could be the main reasons for the tree species richness effect on microbial investment into P acquisition. Unlike N (Parton *et al*., 2007; Hedin *et al*., 2009; Mason *et al*., 2022), P is a growth-limiting element in subtropical forests (Vitousek *et al*., 2010). P is mainly derived from rock weathering, and in natural ecosystems with limited influx or efflux of organic or mineral matter, the P pool is fixed early in development; thus, the P in most natural ecosystems cannot be easily replenished after loss (Vitousek *et al*., 2010). Therefore, P limitation caused by depletion is common in subtropical soils with rapid nutrient turnover and loss (Walker & Syers, 1976). Furthermore, phosphate ions and also enzymes can be sorbed by secondary minerals such as iron and aluminum (oxy)hydroxides in soil (Rojo *et al*., 1990; Burns *et al*., 2013; Nannipieri *et al*., 2018). Therefore, soil microorganisms have to increase the activities of P-acquiring enzymes if they suffer from P limitation and have to satisfy their own demand while potentially competing with plants. The increasing investment in P acquisition with more species-rich tree mixtures also suggests that the P demand is enhanced (Figs. 1f and 2c). Consequently, microorganisms need to invest more energy in the production of nutrient-acquiring enzymes (Sinsabaugh *et al*., 2009; Nannipieri *et al*., 2018). The stoichiometric analysis revealed that tree species richness caused a significant decrease in vector length (Fig. 2, a and b), that is, a decrease in the microbial investment into C relative to nutrient acquisition in litter and mineral soil. This points to another energy supplier, namely the plant. But the additional C source is unlikely to stem primarily from enhanced litter inputs, as such inputs would also stimulate microbial investment into C-acquiring enzymes (Fig. 1a). Instead, it is more likely derived from labile, low-molecular-weight C supplied directly by plant roots. Such readily available C can enter the soil via increased root exudation with higher plant diversity (Steinauer *et al*., 2016; Eisenhauer *et al*., 2017). Supporting this mechanism, ^13^C pulse-labeling experiments have shown that greater plant diversity enhances root-derived C inputs and microbial access to recently fixed C in the rhizosphere, which in turn stimulates soil microbial activity and promotes C support (Lange *et al*., 2015; Mellado-Vázquez *et al*., 2016). These coupled processes may intensify plant–microbe interactions in diverse tree mixtures and provide the additional energy needed to support microbial nutrient acquisition.

### AM-associated tree mixtures drive stronger soil microbial nutrient acquisition

Vector lengths decreased in diverse tree mixtures and particularly in mineral soil of mixtures containing AM-associated trees indicative of less investment into microbial C acquisition (Fig. 4d). Two not mutually exclusive mechanisms may underlie this pattern. First, AM-associated trees may channel a larger proportion of newly assimilated photosynthates belowground, thereby enhancing the supply of C to soil via the root exudate inputs. Recent studies have shown that AM-dominated forests contribute greater root-derived C inputs to soil compared with EcM systems (Keller *et al*., 2021). Research from our study site supports this interpretation (Weinhold *et al*., 2022): coumarins, which are key plant-derived compounds involved in rhizosphere acidification and the mobilization of iron and manganese (Pieterse & Stringlis, 2023), were detected in the root exudates of AM-associated trees (*Cinnamomum camphora* and *Schima superba*), but not in those of EcM-associated trees. Particularly under higher tree diversity and nutrient demand conditions, AM fungi may receive more host photosynthates to sustain nutrient (especially P) acquisition, intensifying C transfer through mycorrhizal pathways as was shown for grassland communities (Lekberg *et al*., 2024). Second, the mycorrhizal type also regulates the quality of root exudation, with AM-associated trees releasing more labile C to the rhizosphere through simple exudates, whereas EcM-associated trees secrete chemically more complex compounds (Liese *et al*., 2017). Consequently, microorganisms in AM-associated tree mixtures may experience higher availability of readily utilizable C, thereby reducing their investment into C acquisition while enhancing their nutrient acquisition. Altogether, this suggests that the interaction between plants and soil microorganisms is intensified by tree species richness, particularly if AM hosts are present.

The host trees of mycorrhizal symbiosis affected the microbial investment into P acquisition: the investment into P acquisition in mineral soil was larger in mixtures with AM- than in those with EcM-associated trees (Fig. 3). Similarly, our vector analysis showed that the investment into P relative to N acquisition gained importance if the AM-associated trees were present as compared to the presence of EcM-associated trees (Fig. 3e). This may be explained by the special nutrient acquisition mode in the AM symbiosis: unlike EcM fungi, AM fungi have a limited ability to mobilize nutrients from SOM, especially to mineralize organic P (Van Der Heijden *et al*., 2008). For instance, the AM fungi lack genes encoding phytases (Tisserant *et al*., 2013), the enzymes required to mineralize phytate, a common but recalcitrant form of organic P in soils (Giaveno *et al*., 2010; Liu *et al*., 2022). Therefore, AM fungi rely on recruiting bacteria with P acquisition function to colonize in their hyphae to enhance the mineralization of organic P, especially monoesters and diesters (Zhang *et al*., 2016). This can be confirmed by the previous studies that found phosphatase activity produced by phosphate-solubilizing bacteria on the hyphae surface of AM fungi (Joner & Johansen, 2000; Feng *et al*., 2002; Ding *et al*., 2021). Furthermore, at our study site, AM-associated microbial subcommunities were more strongly linked to genes encoding P-cycling enzymes than EcM-associated ones (Singavarapu *et al*., 2023). Overall, these mechanisms likely underlie the enhanced microbial investment in P acquisition observed in AM-associated trees, emphasizing the functional complementarity between AM fungi and their associated bacterial partners.

The microbial investment into P acquisition in the presence of AM-associated trees was enhanced with increasing tree species richness (Fig. 4c). The stronger biodiversity–ecosystem functioning relationship in AM-associated tree communities, reflecting higher plant growth and nutrient demand, may also explain their higher nutrient acquisition in enzymatic strategies. This pattern is consistent with evidence from the same experiment site showing that AM-associated trees contribute more strongly to the positive tree diversity–productivity relationship than EcM-associated trees (Deng *et al*., 2023). Additionally, more diverse plants may also increase the biomass and diversity of AM fungi that coexist with roots (Weber *et al*., 2025), which could also enhance the P-acquiring enzyme secretion by soil microorganisms (Mellado-Vázquez *et al*., 2016; Oelmann *et al*., 2021; Qin *et al*., 2022). This highlights the critical role of mycorrhizal fungi and soil microbe–tree interactions in regulating nutrient, especially P, cycling in more diverse tree mixtures.

## Acknowledgements

We thank Wensheng Yang, Zhiyuan Zhang, Hongbo Yang, Aline Ott-Fuchs, Zhigao Li, Sai Peng, Yi Li, and many local assistants for their support in the field work and laboratory work. We are grateful to Yufeng Qiu, Zhuolin Yu, Hui Wang, Yidan Jin, Xiangmei Bao, Na Li, and Wei Liu for their assistance in the laboratory work. We gratefully acknowledge the support of the BEF-China platform and the support of Zhejiang Qianjiangyuan Forest Biodiversity National Observation and Research Station. Our sincere appreciation goes to Keping Ma and Bernhard Schmid for initiating the BEF-China platform. We extend our gratitude to Chao-Dong Zhu for his combined efforts in coordinating the research unit MultiTroph. The study was funded by the Deutsche Forschungsgemeinschaft (DFG, German Research Foundation) – 452861007. Helge Bruelheide also acknowledges support from the International Research Training Group TreeDì, jointly funded by the DFG – 319936945/GRK2324 and the University of the Chinese Academy of Sciences (UCAS). We acknowledge Bo Yang for his assistance in maintaining the research station.

## Author contributions

Y.O. conceived and supported the study and contributed to improving the manuscript. S.S. and T.S. jointly supported the study and also helped revise the draft. N.Z. provided part of the data. Z.Y. participated in data collection. Y.C. provided laboratory facilities. H.B., X.L., A.M.K., and S.L. contributed to the research. H.Z. contributed to study design, data collection, analysis, and manuscript writing. All authors contributed to improving the paper.

## Data availability

The source data of soil and litter enzyme activities, as well as the vector analysis results, will be openly accessible on the BEF-China public data platform via links upon acceptance of the manuscript.

## Supplementary information

Additional Supporting Information may be found online in the Supporting Information section at the end of the article.

**Fig. S1** Litter enzyme activities (a-c) and microbial C vs. nutrient acquisition (d, e) in relation to tree mycorrhizal types for plots with mixtures containing only arbuscular mycorrhizal (AM) host tree species, only ectomycorrhizal (EcM) host tree species, and both mycorrhizal types (AM + EcM).

**Fig. S2** Litter enzyme activities (a-c) and microbial C vs. nutrient acquisition (d-e) in relation to tree species richness (one, two, four) for plots with mixtures containing only arbuscular mycorrhizal (AM) host tree species, only ectomycorrhizal (EcM) host tree species, and both tree mycorrhizal types (AM + EcM).

**Table S1** Linear mixed-effect models for studying soil enzyme activities and microbial C vs. nutrient mobilization as affected by tree species richness (log_2_ scale), tree mycorrhizal types, and topographic factors, with the sampling plot as a random factor.

**Table S2** Linear mixed-effect models for studying litter enzyme activities and microbial C vs. nutrient mobilization as affected by tree species richness (log_2_ scale), tree mycorrhizal types, and topographic factors, with the sampling plot as a random factor.

**Table S3** Linear mixed-effect models for studying soil enzyme activities and microbial C vs. nutrient mobilization as affected by tree species richness (log_2_ scale) and topographic factors in arbuscular mycorrhizal (AM) host plots across a gradient of 1-, 2-, and 4-tree species mixtures, with the sampling plot as a random factor.

**Table S4** Linear mixed-effect models for studying soil enzyme activities and microbial C vs. nutrient mobilization as affected by tree species richness (log_2_ scale) and topographic factors in arbuscular mycorrhizal & ectomycorrhizal (AM + EcM) host plots across a gradient of 1-, 2-, and 4-tree species mixtures, with the sampling plot as a random factor.

**Table S5** Linear mixed-effect models for studying soil enzyme activities and microbial C vs. nutrient mobilization as affected by tree species richness (log_2_ scale) and topographic factors in ectomycorrhizal (EcM) host plots across a gradient of 1-, 2-, and 4-tree species mixtures, with the sampling plot as a random factor.

**Table S6** Linear mixed-effect models for studying litter enzyme activities and microbial C vs. nutrient mobilization as affected by tree species richness (log_2_ scale) and topographic factors in arbuscular mycorrhizal (AM) host plots across a gradient of 1-, 2-, and 4-tree species mixtures, with the sampling plot as a random factor.

**Table S7** Linear mixed-effect models for studying litter enzyme activities and microbial C vs. nutrient mobilization as affected by tree species richness (log_2_ scale) and topographic factors in arbuscular mycorrhizal & ectomycorrhizal (AM + EcM) host plots across a gradient of 1-, 2-, and 4-tree species mixtures, with the sampling plot as a random factor.

**Table S8** Linear mixed-effect models for studying litter enzyme activities and microbial C vs. nutrient mobilization as affected by tree species richness (log_2_ scale) and topographic factors in ectomycorrhizal (EcM) host plots across a gradient of 1-, 2-, and 4-tree species mixtures, with the sampling plot as a random factor.

**Table S9** The 40 local tree species planted in the BEF-China experiment with family and mycorrhizal type and site information.

## References

Bruelheide H, Nadrowski K, Assmann T, Bauhus J, Both S, Buscot F, Chen X-Y, Ding B, Durka W, Erfmeier A, et al. 2014. Designing forest biodiversity experiments: general considerations illustrated by a new large experiment in subtropical China. Methods in Ecology and Evolution 5(1): 74–89.

Burns RG, DeForest JL, Marxsen J, Sinsabaugh RL, Stromberger ME, Wallenstein MD, Weintraub MN, Zoppini A. 2013. Soil enzymes in a changing environment: Current knowledge and future directions. Soil Biology and Biochemistry 58: 216–234.

Čapek P, Manzoni S, Kaštovská E, Wild B, Diáková K, Bárta J, Schnecker J, Biasi C, Martikainen PJ, Alves RJE, et al. 2018. A plant–microbe interaction framework explaining nutrient effects on primary production. Nature Ecology & Evolution 2(10): 1588–1596.

Cardinale BJ, Wright JP, Cadotte MW, Carroll IT, Hector A, Srivastava DS, Loreau M, Weis JJ. 2007. Impacts of plant diversity on biomass production increase through time because of species complementarity. Proceedings of the National Academy of Sciences 104(46): 18123–18128.

Chen X, Chen HYH, Chang SX. 2022. Meta-analysis shows that plant mixtures increase soil phosphorus availability and plant productivity in diverse ecosystems. Nature Ecology & Evolution 6(8): 1112–1121.

Chen X, Chen HYH, Searle EB, Chen C, Reich PB. 2021. Negative to positive shifts in diversity effects on soil nitrogen over time. Nature Sustainability 4(3): 225–232.

Das SK, Varma A 2011. Role of Enzymes in Maintaining Soil Health. In: Shukla G, Varma A eds. Soil Enzymology. Berlin Heidelberg, Germany: Springer Berlin Heidelberg, 25–42.

De Groote SRE, Vanhellemont M, Baeten L, De Schrijver A, Martel A, Bonte D, Lens L, Verheyen K. 2018. Tree species diversity indirectly affects nutrient cycling through the shrub layer and its high-quality litter. Plant and Soil 427(1): 335–350.

Deng M, Hu S, Guo L, Jiang L, Huang Y, Schmid B, Liu C, Chang P, Li S, Liu X, et al. 2023. Tree mycorrhizal association types control biodiversity-productivity relationship in a subtropical forest. Science Advances 9(3): eadd4468.

Ding W, Cong W-F, Lambers H. 2021. Plant phosphorus-acquisition and -use strategies affect soil carbon cycling. Trends in Ecology & Evolution 36(10): 899–906.

Eisenhauer N, Lanoue A, Strecker T, Scheu S, Steinauer K, Thakur MP, Mommer L. 2017. Root biomass and exudates link plant diversity with soil bacterial and fungal biomass. Scientific Reports 7(1): 44641.

Feng G, Su Y, Li X, Wang H, Zhang F, Tang C, Rengel Z. 2002. Histochemical visualization of phosphatase released by arbuscular mycorrhizal fungi in soil. Journal of Plant Nutrition 25(5): 1–1.

Freschet GT, Pagès L, Iversen CM, Comas LH, Rewald B, Roumet C, Klimešová J, Zadworny M, Poorter H, Postma JA, et al. 2021. A starting guide to root ecology: strengthening ecological concepts and standardising root classification, sampling, processing and trait measurements. New Phytologist 232(3): 973–1122.

German DP, Weintraub MN, Grandy AS, Lauber CL, Rinkes ZL, Allison SD. 2011. Optimization of hydrolytic and oxidative enzyme methods for ecosystem studies. Soil Biology and Biochemistry 43(7): 1387–1397.

Giaveno C, Celi L, Richardson AE, Simpson RJ, Barberis E. 2010. Interaction of phytases with minerals and availability of substrate affect the hydrolysis of inositol phosphates. Soil Biology and Biochemistry 42(3): 491–498.

Griffiths HM, Ashton LA, Parr CL, Eggleton P. 2021. The impact of invertebrate decomposers on plants and soil. New Phytologist 231(6): 2142–2149.

Hacker N, Ebeling A, Gessler A, Gleixner G, González Macé O, de Kroon H, Lange M, Mommer L, Eisenhauer N, Ravenek J, et al. 2015. Plant diversity shapes microbe-rhizosphere effects on P mobilisation from organic matter in soil. Ecology Letters 18(12): 1356–1365.

Hedin LO, Brookshire ENJ, Menge DNL, Barron AR. 2009. The Nitrogen Paradox in Tropical Forest Ecosystems. Annual Review of Ecology, Evolution, and Systematics 40(Volume 40, 2009): 613–635.

Hoeber S, Fransson P, Weih M, Manzoni S. 2020. Leaf litter quality coupled to Salix variety drives litter decomposition more than stand diversity or climate. Plant and Soil 453(1): 313–328.

Huang Y, Chen Y, Castro-Izaguirre N, Baruffol M, Brezzi M, Lang A, Li Y, Härdtle W, von Oheimb G, Yang X, et al. 2018. Impacts of species richness on productivity in a large-scale subtropical forest experiment. Science 362(6410): 80–83.

Isbell F, Gonzalez A, Loreau M, Cowles J, Díaz S, Hector A, Mace GM, Wardle DA, O’Connor MI, Duffy JE, et al. 2017. Linking the influence and dependence of people on biodiversity across scales. Nature 546(7656): 65–72.

Jacoby R, Peukert M, Succurro A, Koprivova A, Kopriva S. 2017. The Role of Soil Microorganisms in Plant Mineral Nutrition—Current Knowledge and Future Directions. Frontiers in Plant Science 8.

Jiang F, Zhang L, Zhou J, George TS, Feng G. 2021. Arbuscular mycorrhizal fungi enhance mineralisation of organic phosphorus by carrying bacteria along their extraradical hyphae. New Phytologist 230(1): 304–315.

Joner EJ, Johansen A. 2000. Phosphatase activity of external hyphae of two arbuscular mycorrhizal fungi. Mycological Research 104(1): 81–86.

Keller AB, Brzostek ER, Craig ME, Fisher JB, Phillips RP. 2021. Root-derived inputs are major contributors to soil carbon in temperate forests, but vary by mycorrhizal type. Ecology Letters 24(4): 626–635.

Lambers H, Albornoz F, Kotula L, Laliberté E, Ranathunge K, Teste FP, Zemunik G. 2018. How belowground interactions contribute to the coexistence of mycorrhizal and non-mycorrhizal species in severely phosphorus-impoverished hyperdiverse ecosystems. Plant and Soil 424(1): 11–33.

Lange M, Eisenhauer N, Sierra CA, Bessler H, Engels C, Griffiths RI, Mellado-Vázquez PG, Malik AA, Roy J, Scheu S, et al. 2015. Plant diversity increases soil microbial activity and soil carbon storage. Nature Communications 6(1): 6707.

Lekberg Y, Jansa J, McLeod M, DuPre ME, Holben WE, Johnson D, Koide RT, Shaw A, Zabinski C, Aldrich-Wolfe L. 2024. Carbon and phosphorus exchange rates in arbuscular mycorrhizas depend on environmental context and differ among co-occurring plants. New Phytologist 242(4): 1576–1588.

Li Y, Bruelheide H, Scholten T, Schmid B, Sun Z, Zhang N, Bu W, Liu X, Ma K. 2019. Early positive effects of tree species richness on soil organic carbon accumulation in a large-scale forest biodiversity experiment. Journal of Plant Ecology 12(5): 882–893.

Liese R, Lübbe T, Albers NW, Meier IC. 2017. The mycorrhizal type governs root exudation and nitrogen uptake of temperate tree species. Tree Physiology 38(1): 83–95.

Lindahl BD, Tunlid A. 2015. Ectomycorrhizal fungi – potential organic matter decomposers, yet not saprotrophs. New Phytologist 205(4): 1443–1447.

Liu X, Han R, Cao Y, Turner BL, Ma LQ. 2022. Enhancing Phytate Availability in Soils and Phytate-P Acquisition by Plants: A Review. Environmental Science & Technology 56(13): 9196–9219.

Martin F, Aerts A, Ahrén D, Brun A, Danchin EGJ, Duchaussoy F, Gibon J, Kohler A, Lindquist E, Pereda V, et al. 2008. The genome of Laccaria bicolor provides insights into mycorrhizal symbiosis. Nature 452(7183): 88–92.

Mason RE, Craine JM, Lany NK, Jonard M, Ollinger SV, Groffman PM, Fulweiler RW, Angerer J, Read QD, Reich PB, et al. 2022. Evidence, causes, and consequences of declining nitrogen availability in terrestrial ecosystems. Science 376(6590): eabh3767.

Mellado-Vázquez PG, Lange M, Bachmann D, Gockele A, Karlowsky S, Milcu A, Piel C, Roscher C, Roy J, Gleixner G. 2016. Plant diversity generates enhanced soil microbial access to recently photosynthesized carbon in the rhizosphere. Soil Biology and Biochemistry 94: 122–132.

Moorhead DL, Sinsabaugh RL, Hill BH, Weintraub MN. 2016. Vector analysis of ecoenzyme activities reveal constraints on coupled C, N and P dynamics. Soil Biology and Biochemistry 93: 1–7.

Nannipieri P, Giagnoni L, Renella G, Puglisi E, Ceccanti B, Masciandaro G, Fornasier F, Moscatelli MC, Marinari S. 2012. Soil enzymology: classical and molecular approaches. Biology and Fertility of Soils 48(7): 743–762.

Nannipieri P, Trasar-Cepeda C, Dick RP. 2018. Soil enzyme activity: a brief history and biochemistry as a basis for appropriate interpretations and meta-analysis. Biology and Fertility of Soils 54(1): 11–19.

Oelmann Y, Lange M, Leimer S, Roscher C, Aburto F, Alt F, Bange N, Berner D, Boch S, Boeddinghaus RS, et al. 2021. Above- and belowground biodiversity jointly tighten the P cycle in agricultural grasslands. Nature Communications 12(1): 4431.

Oelmann Y, Richter AK, Roscher C, Rosenkranz S, Temperton VM, Weisser WW, Wilcke W. 2011. Does plant diversity influence phosphorus cycling in experimental grasslands? Geoderma 167–168: 178-187.

Parton W, Silver WL, Burke IC, Grassens L, Harmon ME, Currie WS, King JY, Adair EC, Brandt LA, Hart SC, et al. 2007. Global-Scale Similarities in Nitrogen Release Patterns During Long-Term Decomposition. Science 315(5810): 361–364.

Pieterse CMJ, Stringlis IA. 2023. Chemical symphony of coumarins and phenazines in rhizosphere iron solubilization. Proceedings of the National Academy of Sciences 120(18): e2304171120.

Pritsch K, Garbaye J. 2011. Enzyme secretion by ECM fungi and exploitation of mineral nutrients from soil organic matter. Annals of Forest Science 68(1): 25–32.

Qin Y, Zhang W, Feng Z, Feng G, Zhu H, Yao Q. 2022. Arbuscular mycorrhizal fungus differentially regulates P mobilizing bacterial community and abundance in rhizosphere and hyphosphere. Applied Soil Ecology 170: 104294.

Rojo MJ, Carcedo SG, Mateos MP. 1990. Distribution and characterization of phosphatase and organic phosphorus in soil fractions. Soil Biology and Biochemistry 22(2): 169–174.

Rosling A, Midgley MG, Cheeke T, Urbina H, Fransson P, Phillips RP. 2016. Phosphorus cycling in deciduous forest soil differs between stands dominated by ecto- and arbuscular mycorrhizal trees. New Phytologist 209(3): 1184–1195.

Scherber C, Eisenhauer N, Weisser WW, Schmid B, Voigt W, Fischer M, Schulze E-D, Roscher C, Weigelt A, Allan E, et al. 2010. Bottom-up effects of plant diversity on multitrophic interactions in a biodiversity experiment. Nature 468(7323): 553–556.

Schnitzer SA, Klironomos JN, HilleRisLambers J, Kinkel LL, Reich PB, Xiao K, Rillig MC, Sikes BA, Callaway RM, Mangan SA, et al. 2011. Soil microbes drive the classic plant diversity–productivity pattern. Ecology 92(2): 296–303.

Scholten T, Goebes P, Kühn P, Seitz S, Assmann T, Bauhus J, Bruelheide H, Buscot F, Erfmeier A, Fischer M, et al. 2017. On the combined effect of soil fertility and topography on tree growth in subtropical forest ecosystems—a study from SE China. Journal of Plant Ecology 10(1): 111–127.

Seitz S, Goebes P, Song Z, Bruelheide H, Härdtle W, Kühn P, Li Y, Scholten T. 2016. Tree species and functional traits but not species richness affect interrill erosion processes in young subtropical forests. SOIL 2(1): 49–61.

Semchenko M, Barry KE, de Vries FT, Mommer L, Moora M, Maciá-Vicente JG. 2022. Deciphering the role of specialist and generalist plant–microbial interactions as drivers of plant–soil feedback. New Phytologist 234(6): 1929–1944.

Singavarapu B, Du J, Beugnon R, Cesarz S, Eisenhauer N, Xue K, Wang Y, Bruelheide H, Wubet T. 2023. Functional Potential of Soil Microbial Communities and Their Subcommunities Varies with Tree Mycorrhizal Type and Tree Diversity. Microbiology Spectrum 11(2): e04578–04522.

Sinsabaugh RL, Hill BH, Follstad Shah JJ. 2009. Ecoenzymatic stoichiometry of microbial organic nutrient acquisition in soil and sediment. Nature 462(7274): 795–798.

Smith SE, Read DJ. 2008. Mycorrhizal symbiosis. 3rd edn. Academic Press, London, UK.

Smith SE, Smith FA. 2011. Roles of Arbuscular Mycorrhizas in Plant Nutrition and Growth: New Paradigms from Cellular to Ecosystem Scales. Annual Review of Plant Biology 62(Volume 62, 2011): 227–250.

Steidinger BS, Crowther TW, Liang J, Van Nuland ME, Werner GDA, Reich PB, Nabuurs GJ, de-Miguel S, Zhou M, Picard N, et al. 2019. Climatic controls of decomposition drive the global biogeography of forest-tree symbioses. Nature 569(7756): 404–408.

Steinauer K, Chatzinotas A, Eisenhauer N. 2016. Root exudate cocktails: the link between plant diversity and soil microorganisms? Ecology and Evolution 6(20): 7387–7396.

Talbot JM, Bruns TD, Smith DP, Branco S, Glassman SI, Erlandson S, Vilgalys R, Peay KG. 2013. Independent roles of ectomycorrhizal and saprotrophic communities in soil organic matter decomposition. Soil Biology and Biochemistry 57: 282–291.

Tedersoo L, Bahram M. 2019. Mycorrhizal types differ in ecophysiology and alter plant nutrition and soil processes. Biological Reviews 94(5): 1857–1880.

Tedersoo L, Bahram M, Zobel M. 2020. How mycorrhizal associations drive plant population and community biology. Science 367(6480): eaba1223.

Tisserant E, Malbreil M, Kuo A, Kohler A, Symeonidi A, Balestrini R, Charron P, Duensing N, Frei dit Frey N, Gianinazzi-Pearson V, et al. 2013. Genome of an arbuscular mycorrhizal fungus provides insight into the oldest plant symbiosis. Proceedings of the National Academy of Sciences 110(50): 20117–20122.

Van Der Heijden MGA, Bardgett RD, Van Straalen NM. 2008. The unseen majority: soil microbes as drivers of plant diversity and productivity in terrestrial ecosystems. Ecology Letters 11(3): 296–310.

Vitousek PM, Porder S, Houlton BZ, Chadwick OA. 2010. Terrestrial phosphorus limitation: mechanisms, implications, and nitrogen–phosphorus interactions. Ecological Applications 20(1): 5–15.

Wagg C, Roscher C, Weigelt A, Vogel A, Ebeling A, de Luca E, Roeder A, Kleinspehn C, Temperton VM, Meyer ST, et al. 2022. Biodiversity–stability relationships strengthen over time in a long-term grassland experiment. Nature Communications 13(1): 7752.

Walker TW, Syers JK. 1976. The fate of phosphorus during pedogenesis. Geoderma 15(1): 1–19.

Wang G, Jin Z, George TS, Feng G, Zhang L. 2023. Arbuscular mycorrhizal fungi enhance plant phosphorus uptake through stimulating hyphosphere soil microbiome functional profiles for phosphorus turnover. New Phytologist 238(6): 2578–2593.

Weber SE, Bascompte J, Kahmen A, Niklaus PA. 2025. AMF diversity promotes plant community phosphorus acquisition and reduces carbon costs per unit of phosphorus. New Phytologist 248(2): 886–896.

Weinhold A, Döll S, Liu M, Schedl A, Pöschl Y, Xu X, Neumann S, van Dam NM. 2022. Tree species richness differentially affects the chemical composition of leaves, roots and root exudates in four subtropical tree species. Journal of Ecology 110(1): 97–116.

Yang X, Bauhus J, Both S, Fang T, Härdtle W, Kröber W, Ma K, Nadrowski K, Pei K, Scherer-Lorenzen M, et al. 2013. Establishment success in a forest biodiversity and ecosystem functioning experiment in subtropical China (BEF-China). European Journal Of Forest Research 132(4): 593–606.

Zhang H, Fang Y, Zhang B, Luo Y, Yi X, Wu J, Chen Y, Sarker TC, Cai Y, Chang SX. 2022. Land-use-driven change in soil labile carbon affects microbial community composition and function. Geoderma 426: 116056.

Zhang L, Xu M, Liu Y, Zhang F, Hodge A, Feng G. 2016. Carbon and phosphorus exchange may enable cooperation between an arbuscular mycorrhizal fungus and a phosphate-solubilizing bacterium. New Phytologist 210(3): 1022–1032.

Zhang N, Li Y, Wubet T, Bruelheide H, Liang Y, Purahong W, Buscot F, Ma K. 2018. Tree species richness and fungi in freshly fallen leaf litter: Unique patterns of fungal species composition and their implications for enzymatic decomposition. Soil Biology and Biochemistry 127: 120–126.

